# Simultaneous immunomodulation and epithelial-to-mesenchymal transition drives lung adenocarcinoma progression

**DOI:** 10.1101/2025.02.19.637138

**Authors:** Junbum Kim, Hiranmayi Ravichandran, Liron Yoffe, Bhavneet Bhinder, Kyle Finos, Arshdeep Singh, Bradley B Pua, Stewart Bates, Bevan Emma Huang, Andre F. Rendeiro, Vivek Mittal, Nasser K. Altorki, Timothy E. McGraw, Olivier Elemento

## Abstract

Lung cancer remains the deadliest cancer in the United States, with lung adenocarcinoma (LUAD) as its most prevalent subtype. While computed tomography (CT)-based screening has improved early detection and enabled curative surgeries, the molecular and cellular dynamics driving early-stage LUAD progression remain poorly understood, limiting non-surgical treatment options. To address this gap, we profiled 2.24 million cells from 122 early-stage LUAD patients using multiplexed imaging mass cytometry (IMC). This analysis revealed the molecular, spatial, and temporal dynamics of LUAD development.

Our findings uncover a binary progression model. LUAD advances through either inflammation, driven by a balance of cytotoxic and regulatory immune activity, or fibrosis, characterized by stromal activation. Surprisingly, tumor cell populations did not increase significantly. Instead, they displayed a mixed phenotypic profile consistent with epithelial-to-mesenchymal transition (EMT), effectively masking the expansion of malignant cells.

Furthermore, we addressed discrepancies between CT-based and histology-based subtyping. CT scans, while non-invasive, often mischaracterize invasive fibrotic tumors—which account for 20.5% of LUAD cases—as mild, non-solid ground glass opacities (GGOs). Using high-content IMC imaging, we demonstrate that these tumors harbor significant risks and advocate for improved diagnostic strategies. These strategies should integrate molecular profiling to refine patient stratification and therapeutic decision-making.

Altogether, our study provides a high-resolution, systems-level view of the tumor microenvironment in early-stage LUAD. We characterize key transitions in oncogenesis and propose a precision-driven framework to enhance the detection and management of aggressive disease subtypes.

## Introduction

Lung cancer ranks among the most prevalent and lethal cancers in the United States, with a five-year survival rate of 20-25%^1–3^. Lung adenocarcinoma (LUAD) is the most commonly observed primary lung cancer in the country, representing about one-half of lung cancer cases^4^. Approximately 74% of lung cancer are diagnosed at advanced stages, where the tumor has already metastasized to regional or distant sites, limiting treatment options. To address this issue, clinical guidelines^5,6^ in the last decade have called for lung cancer screening with CT in high-risk individuals. This has enabled the detection of lung cancer at earlier stages than before, resulting in curative surgeries for lesions exhibiting malignant or premalignant characteristics. However, more detailed cellular evolution and spatial rearrangements underlying the oncogenesis and progression of early stage LUAD^7–9^ remains unclear.

To date, researchers have been trying to identify genomic and proteomic alterations that fosters a protumorigenic microenvironment that allow for LUAD to initiate and progress. Our group previously shared results demonstrating the immune and stromal evolution of the tumor microenvironment through whole exome sequencing and bulk RNA sequencing^10^. Some of these immune adaptations include elevated cytokine expression, CD8 T cell mediated cytotoxicity, and immunosuppression by regulatory T cells. In addition, we previously showed an increase in extracellular matrix (ECM) modifying enzymes and fibroblast disorganization as key alterations.

Recent advancements in technologies like multiplexed imaging^11–14^ and spatial transcriptomics^15–22^ have revolutionized the observation of cellular phenotypes and interactions within native tissue environments, significantly enhancing our understanding of tumors within their stromal and immune contexts. Despite these technological advancements, no studies to date have dedicated an investigation into the detailed molecular and spatial alterations associated with the progression of early-stage LUAD^23–27^ using leveraging technology.

In this study, we seek to further elucidate the spatiomolecular landscape of early-stage LUAD. Our goal is to bridge how oncogenic changes manifest at cellular, structural, and at patient level through imaging mass cytometry (IMC). We specifically look to investigate how cytotoxic and regulatory immune activity can simultaneously drive tumorigenesis along with stromal modification.

## Results

To study molecular changes associated with tumor progression, we collected surgically resected lesions from 122 early-stage LUAD patients (**Figure 1a-b, Extended Figure 1a**). Most patients were diagnosed with stage 1 or 2 LUAD, with a median tumor size of 1.4 cm (**Extended Figure 1b**). Detailed demographic information for the patients are described in **Extended Figure 1b-e**. These specimens were independently graded by expert pathologists and radiologists. The tissue samples were categorized into 44 AIS (adenocarcinoma in situ), 44 MIA (minimally invasive adenocarcinoma), and 35 IAC (invasive adenocarcinoma) cases (**Figure 1b**) histologically. Radiographically, the lesions were classified as 24 solid (S), 31 part-solid (PS), and 62 pure ground-glass nodules (PNS) (**Figure 1b**).

**Figure 1:**
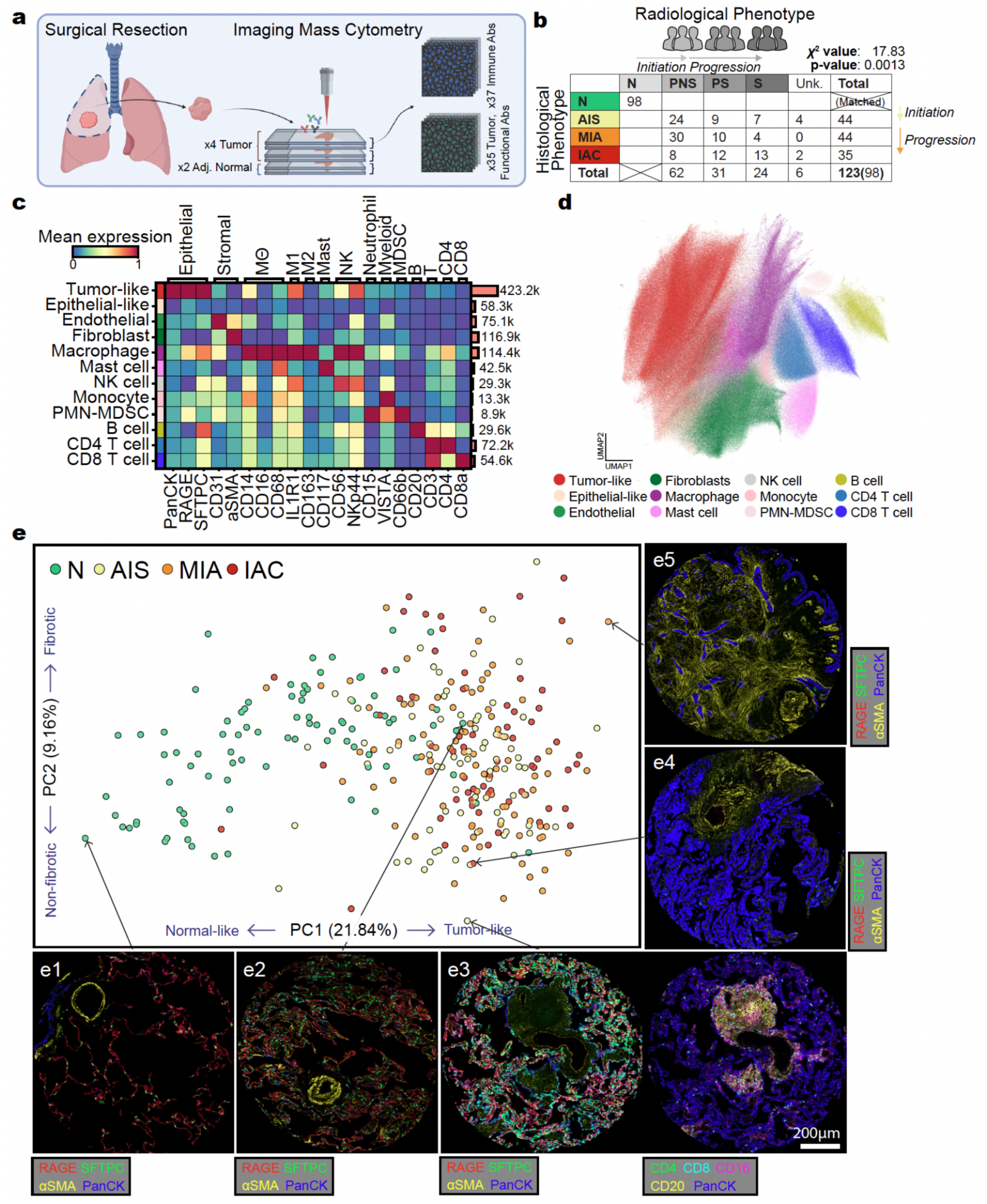
Study design and data overview. **a**) Schematic of study sample collection using surgically resected cancer lesions. Two tumors and one adjacent normal region of interest (ROI) of each patient were selected. Each ROI was imaged twice, using adjacent sections. Each section was stained with a tumor- and immune-specific cocktail of antibodies. **b**) Composition of the cohort stratified by histological and radiological classifications. Histological groups include adenocarcinoma in-situ (AIS), minimally invasive adenocarcinoma (MIA), and invasive adenocarcinoma (IAC). Radiological groups were pure non-solid (PNS), part-solid (PS), and solid (S). **c**) Heatmap of identified cell phenotypes. Colors represent mean expression profiles in the tumor characterization panel. Bar plot on the right indicates the number of cells for each cell phenotype. **d**) Uniform manifold approximation and projection (UMAP) of cell types for the tumor characterization panel. **e**) Principal component analysis plot in the middle of the figure visualizes general molecular trends and variation across ROIs. Each sample in the plot represents cell type density of two adjacent ROI stacks and are colored by histology subtype. Surrounding images visualize the epithelial, stromal, and immune molecular characteristics of selected ROIs.

For each patient, we selected up to two tumor samples and one adjacent-normal region of interest (ROI). We prepared two tissue slides along the z-stack for Imaging Mass Cytometry (IMC) to analyze both the tumor and immune characteristics (**Data Table 1, Extended Figure 2**) for selected ROIs. To characterize the tumor evolution, we defined disease initiation as the transition from adjacent-normal to AIS, and tumor progression as the shift from AIS to more advanced stages like MIA and IAC (**Figure 1b**). To extensively characterize normal lung behavior, which serves as an important anchor-point for our experiment, we collected a total of 98 matched adjacent-normal tissues from our patients (**Figure 1b**).

**Data Table 1.** List of Antibodies used for Imaging Mass Cytometry data acquisition.

| Channel | Metal | Tumor Panel | Immune Panel |
| --- | --- | --- | --- |
| 1 | Pr141 | HLADR | HLADR |
| 2 | Nd142 | NKp44 | GranzymeB |
| 3 | Nd143 | CD31 | Vimentin |
| 4 | Nd144 | CD14 | CD14 |
| 5 | Nd145 | CD33 | Tbet |
| 6 | Nd146 | αSMA | CD16 |
| 7 | Sm147 | CD163 | CD163 |
| 8 | Nd148 | PanCytokeratin | ICOS |
| 9 | Sm149 | IL23R | CCR4 |
| 10 | Nd150 | CD103 | PDL1 |
| 11 | Eu151 | CD16 | CD31 |
| 12 | Sm152 | CD66b | CD45 |
| 13 | Eu153 | IL1R1 | LAG3 |
| 14 | Sm154 | IL23p19 | CD11c |
| 15 | Gd155 | RAGE | FoxP3 |
| 16 | Gd156 | CD4 | CD4 |
| 17 | Gd158 | CD15 | CCR7 |
| 18 | Tb159 | CD68 | CD68 |
| 19 | Gd160 | IFNg | CD28 |
| 20 | Dy161 | CD20 | CD20 |
| 21 | Dy162 | CD8a | CD8a |
| 22 | Dy163 | SCL5A2SGLT2 | CD56 |
| 23 | Dy164 | CD117 | CD11b |
| 24 | Ho165 | IL12p40 | PD1 |
| 25 | Er166 | IL1beta | CD45RA |
| 26 | Er167 | HLAABC | CD206 |
| 27 | Er168 | SFTPC | Ki67 |
| 28 | Tm169 | CD56 | CD57 |
| 29 | Er170 | CD3 | CD3 |
| 30 | Yb171 | GLUT5 | CD27 |
| 31 | Yb172 | IL1alpha | 41BB |
| 32 | Yb173 | VISTA | CD45RO |
| 33 | Yb174 | IL17A | TIM3 |
| 34 | Lu175 | FOLR1 | CD25 |
| 35 | Yb176 | MMP7 | CTLA4 |
| 36 | BCKG190 | EMPTY | EMPTY |
| 37 | Ir191 | DNA1 | DNA1 |
| 38 | Ir193 | DNA2 | DNA2 |
| 39 | Pt196 | EMPTY | αSMA |
| 40 | Pt198 | EMPTY | PanCytokeratin |

**Figure 2:**
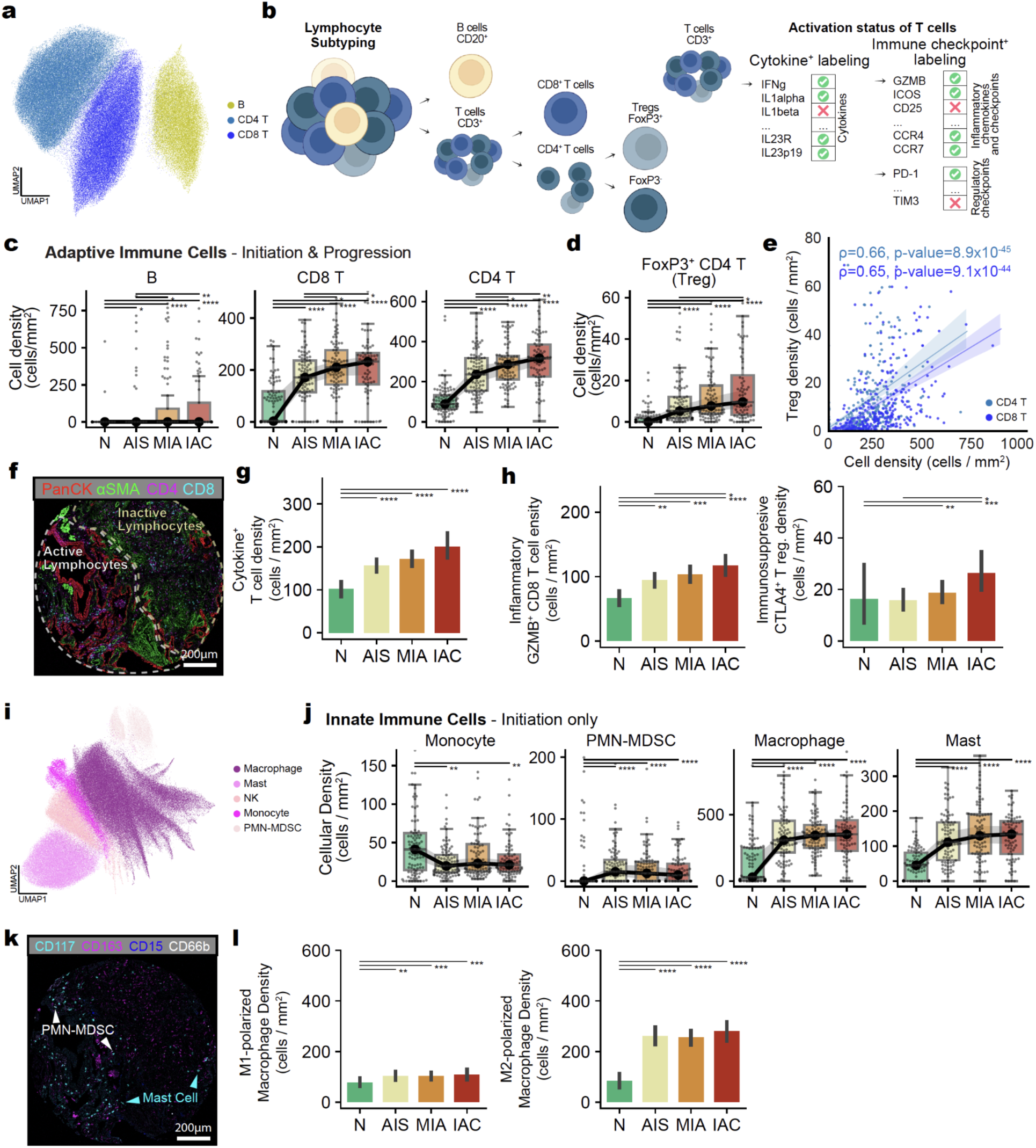
Characterization of immune cells in early-stage LUAD. **a**) UMAP of lymphocytes found in the dataset. **b**) Schematic diagram illustrating the detailed T cell subtype labeling. CD3+ T cells were stratified by FoxP3, CD4, and CD8 expression. All T cells were also annotated for whether they were positive for cytokines and immune checkpoints. **c**) Changes in lymphocyte density represented by boxplots. Line plots were overlaid to highlight the median trends. **d**) Change in Tregs, a subset of CD4 T cell, density. **e**) Scatterplot showing the Spearman’s correlation between Tregs and lymphocytes in each ROI. Tregs were more common in lymphocyte infiltrated ROIs. **f**) Illustrative image of a T cell enriched ROI. PanCK is in red, αSMA is in green, CD4 is in magenta, and CD8 is in cyan. Active lymphocytes were populated towards the bottom left of the ROI where the tumor cells are. **g-h**) Barplot showing change in Cytokine+ T cells, GZMB+ CD8 T cells, and CTLA4+ Treg density across LUAD subtypes. **i**) UMAP of myeloid cells found in the dataset. **j**) Changes in myeloid cell density represented by boxplots. **k**) Illustrative image of myeloid cells in the same ROI as in panel **f**. CD117 is in cyan, CD163 is in magenta, CD15 is in blue, and CD66b is in white. **l**) Barplot showing change in M1 and M2 polarization of macrophages across LUAD subtypes. For **c-d,g-h,j,l**) stars represent statistically significant p-values for two-sided Mann-Whitney U-test with BH multiple hypothesis correction. * p<0.05, ** p<0.01, *** p<0.001, **** p<0.0001.

Using single-cell resolution spatial imaging, we profiled a diverse and well-balanced population of cells from epithelial, stromal, and immune origins. In total, we analyzed 2.24 million cells across two IMC panels (**Figure 1c-d, Extended Figure 3a-d**), including 839.5k epithelial-like cells that ranged from normal lung epithelial cells to tumor cells. We also identified large numbers of tumor-infiltrating immune cells, including 213.4k macrophages, 75.6k B cells, 117.8k CD8 T cells, 163.0k CD4 T cells, 4.2k Tregs (regulatory T cells), 66.7k NK cells, 13.3k neutrophils, and 8.9k polymorphonuclear myeloid-derived suppressor cells (PMN-MDSCs). Among the stromal populations, we found 223.4k fibroblasts, 147.9k endothelial cells, and 128.7k mesenchymal cells (**Extended Figure 3**).

**Figure 3:**
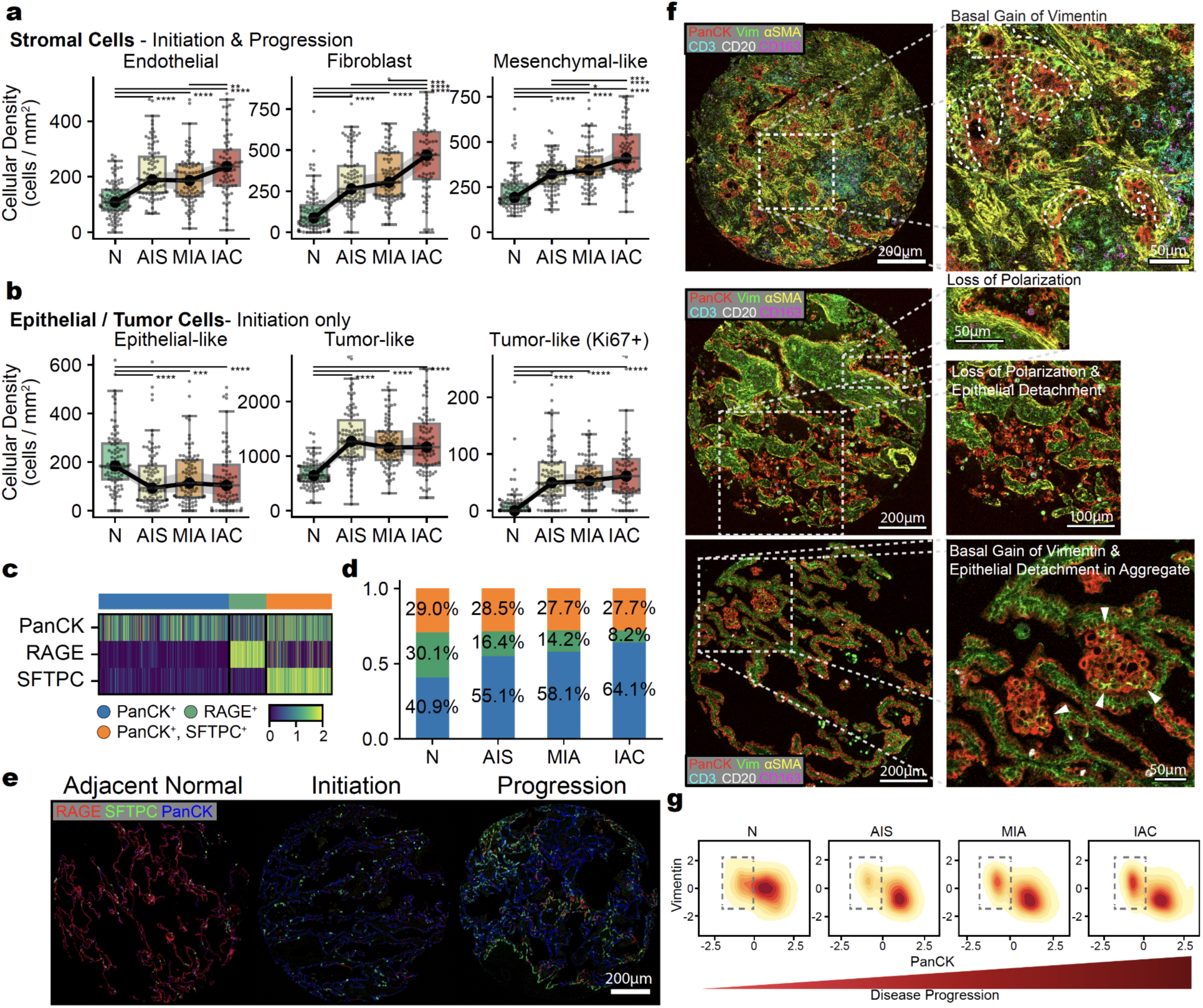
Characterization of stromal, epithelial, and tumor cells in early-stage LUAD. **a-b**) Changes in cell density across histopathology grading represented by boxplots. Line plots highlight the trends. **a**) Change in stromal cell density. **b**) Change in epithelial/tumor cell density. **c**) Heatmap illustrating expression of PanCytokeratin, RAGE, and SFTPC on cells of epithelial origin. **d**) Proportion of three cell subtypes across histopathological subtypes. **e**) Visual epithelial/tumor cell characterization based on PanCytokeratin, RAGE, and SFTPC for adjacent normal, initiation, and progression. **f**) Visualization of ongoing epithelial-to-mesenchymal transition. First image demonstrates the gain of vimentin expression in the basal membrane of epithelial cells. Second image visualizes loss of polarization and epithelial detachment. Third image shows basal gain of vimentin and epithelial detachment in groups of epithelial cells. Scale bars represent 200 µm. **g**) Kernel density estimator plots demonstrating shifts from epithelial to mesenchymal transitions of epithelial cells in more solid GGOs and in more invasive histological subtypes. Increase in PanCK-Vim+ subtype is highlighted in gray box.

Leveraging the cellular phenotypes, we explored the molecular variation underlying the early-stage LUAD cohort. To do this, we performed a principal component analysis (PCA) using the combined cell type densities from the two IMC panels. We highlighted several regions of interest (ROIs) to showcase key features of early-stage LUAD compared to adjacent-normals (**Figure 1e1-5**). Unsurprisingly, the main separation in the data was between adjacent-normal and tumor ROIs. Adjacent-normal samples clustered toward the negative PC1 axis, while relatively advanced tumor subtypes were found on the positive side.

For example, **Figure 1e1** represented the “most normal” ROI, featuring a balanced mixture of AT1 and AT2 cells. This region contained mostly RAGE+ AT1 cells and a smaller number of SFTPC+ AT2 cells. In contrast, some adjacent-normal samples, located closer to the positive PC1 axis, displayed more “tumor-like” features. In one such ROI (**Figure 1e2**), we saw signs of hyperplasia and an increased density of SFTPC+ AT2 cells, commonly thought to be the cells of origin for LUAD. ROIs from AIS samples (**Figure 1e3**) showed an even higher abundance of SFTPC+ AT2 cells. Many of these SFTPC+ cells were also positive for PanCK in tumor samples. As the disease progressed to more invasive stages, ROIs (**Figure 1e4**) revealed tumor cells that were exclusively PanCK+, without the expression of RAGE or SFTPC.

In addition to the contrast between epithelial cells (**Figure 1e1-4**), other significant variations included differences in immune cell composition and the degree of fibrosis. For instance, the AIS ROI in **Figure 1e3** contained two large tertiary lymphocyte structures (TLS) along with smaller lymphocyte clusters, suggesting a high level of immune cell infiltration. **Figure 1e5** highlighted tissue with extreme fibrosis.

**Figure 4:**
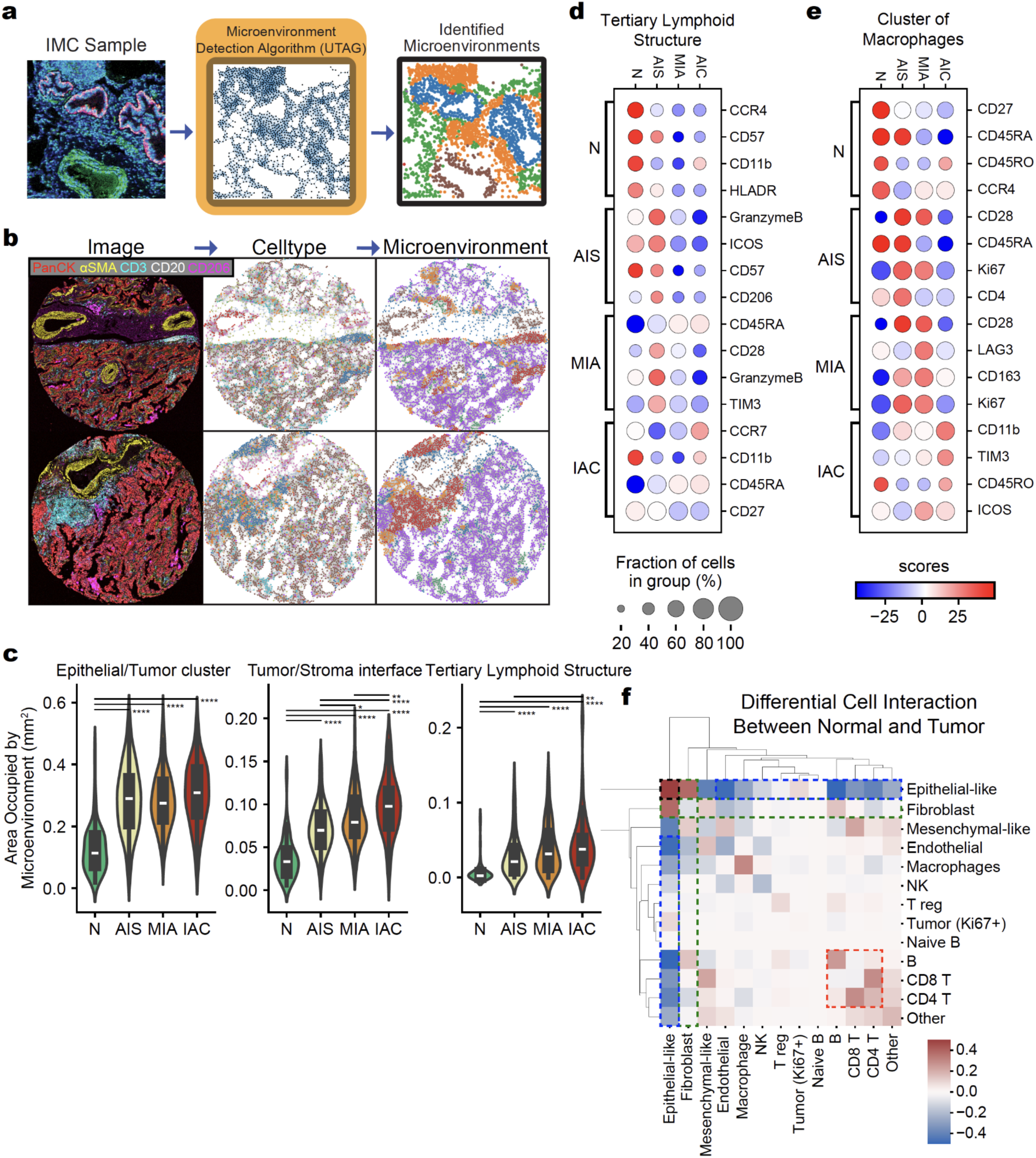
Dynamics of microenvironment and cellular interactions in early-stage LUAD. **a**) Illustrative schematic for niche detection algorithm UTAG. **b**) Descriptive images visualizing the captured cell types and microenvironments through UTAG. **c**) Statistical comparison of area occupied by microenvironment across histopathology subgroups. * p<0.05, ** p<0.01, *** p<0.001, **** p<0.0001, two-sided Mann-Whitney U-test with BH multiple hypothesis correction. **d**) Dotplot visualizing functional shift in tertiary lymphoid structures. **e**) Dotplot visualizing functional shift in clusters of macrophages. **d-e**) Sizes of the dots represent the fraction of cells the marker is expressed in. **f**) Differential cellular interaction between solid and adjacent normal GGO. Positive interaction indicates relative proximity and negative interaction represents relative avoidance. Black dotted box represents epithelial-epithelial interaction. Blue dotted box represents epithelial-other interactions. Green dotted box represents fibroblast interactions. Red dotted box represents interactions from tertiary lymphoid structure.

We applied PCA to illustrate key variations in the LUAD microenvironments captured by our IMC dataset. We qualitatively observed that some of these changes, such as the shift from PanCK- to PanCK+ epithelial cells, were linked to disease progression. With these insights, our next step was to systematically investigate how the variations in immune, stromal, and tumor cells were associated with LUAD progression.

### Simultaneous activation and regulation of immune system drives early-stage LUAD progression

Our explorations through PCA led us to hypothesize a multistep model for tumor development, involving changes in the tumor, stromal, and immune cells. We predicted that there would be an early and significant depletion of the innate immune cells, followed by a gradual exhaustion of the adaptive immune cells. We propose that this sequential shift creates an increasingly pro-inflammatory and immunosuppressive microenvironment.

To test this hypothesis, we first measured the abundance of lymphocytes at different stages of the disease (**Figure 2a**). For detailed comparison, we broke down our lymphocytes to B cells and T cells (**Figure 2b**), categorized our T cells into CD4 T cells and CD8 T cells, and filtered out FoxP3+ CD4 T cells as regulatory T cells (Tregs). In doing so, we identified a synchronous increase in B cells, CD4 T cells, and CD8 T cells (**Figure 2c**). The sharpest increase occurred during the transition from adjacent-normal tissue to tumor tissue with an adjusted p-value as high as 1.78 x 10^-11^. With progression, these cells gradually change from AIS to IAC with a maximum p-value of 0.03 across the tumors. These results suggest a surge in immune activity during the early stages of tumor initiation, which continues to escalate as the disease progresses.

At the same time, we observed an increase in FoxP3+ Tregs (**Figure 2d**). Specifically, FoxP3+ Tregs increase was associated with cytotoxic CD8 T cell and CD4 T helper cell elevation (**Figure 2e**), with a Spearman’s correlation of 0.65 and 0.66 respectively. This suggested that as the tumor progressed, lymphocytes maintained a balance between cytotoxic and regulatory T cell counts, likely contributing to a heightened but regulated inflammatory response.

Given the simultaneous rise in cytotoxic and regulatory T cell counts, we hypothesized that their antagonistic functions would be elevated as well. To test this, we focused on specific T cell subpopulations such as, cytokine-exposed T cells, GZMB+ CD8 T cells, and CTLA4+ Tregs (**Figure 2g-h**). As we expected, we found that the density of cytokine-exposed T cells increased as LUAD progressed (**Figure 2g**). Similarly, we observed an enrichment of both GZMB+ CD8 T cells and CTLA4+ Tregs (**Figure 2h**), confirming that the functional activity of both cytotoxic and regulatory T cells was elevated during disease progression.

In contrast to adaptive immune cells, we hypothesized that innate immune cells would have a limited capacity to respond to tumor progression, as they lack adjustability. Our results confirmed this hypothesis, showing abrupt but restricted changes in the innate immune cells. We observed an increase in monocytes, mast cells, macrophages, and CD15+ CD66b+ polymorphonuclear myeloid-derived suppressor cells (PMN-MDSCs) (**Figure 2m**) exclusively at initiation. Nevertheless, these cells did not change in population with disease progression.

Similar to adaptive immune cells, these cells displayed both pro- and anti-inflammatory characteristics. For instance, we observed an increase in inflammatory cells, such as CD117+ mast cells (**Figure 2k**) and M1 macrophages (**Figure 2l**), which are known to promote inflammation. At the same time, we also saw a significant rise in anti-inflammatory cells, including PMN-MDSCs (**Figure 2k**) and M2 macrophages (**Figure 2l**).

Integrating these findings, we observed a multistep immune modulation during the initiation and progression of LUAD. During initiation, there was a marked shift detected across all immune cell types. As disease progressed, innate immune cells exhibited a diminished or inadequate response, whereas adaptive immune cells continued to display activity. Notably, we report a concurrent increase in cytotoxic and regulatory T cells throughout the course of LUAD progression, resulting in an inflammatory pro-oncogenic immune microenvironment.

### EMT drives LUAD progression

Moving beyond the immune system, we next aimed to investigate how stromal and epithelial cells were affected by LUAD.

In stromal cells, we observed a significant increase in several cell populations. CD31+ endothelial cells, αSMA+ fibroblasts, and Vim+ mesenchymal cells increased with a maximum adjusted p-value of 0.046 (**Figure 3a**). These findings suggested active angiogenesis, fibrosis, and wound healing in the tissue. Interestingly, these trends not only persisted but also intensified as the tumor progressed, indicating a continuous remodeling of the stromal environment throughout tumor development.

In contrast, the epithelial compartment exhibited unexpected dynamics. Although the transition from adjacent normal to tumor tissue was marked by a significant shift, epithelial cells did not demonstrate a consistent increase in abundance throughout tumor progression (**Figure 3b**). Specifically, we did not observe a significant increase in tumor cell density in early-stage LUADs. This finding was unexpected, particularly given the pronounced changes observed in immune and stromal cell populations during the same stages.

Despite the lack of a significant increase in tumor cell numbers during LUAD progression, we sought to determine whether molecular changes occurred within the epithelial compartment. Our analysis revealed a decrease in PanCK-epithelial-like cells, accompanied by an increase in PanCK+ tumor-like cells (**Figure 3b**). Among the tumor-like cells, there was a non-significant rise in the Ki67+ proliferative subset. These cells were further classified into three distinct populations: PanCK+, RAGE+, and PanCK+SFTPC+ (**Figure 3c**). Along the progression axis, we observed a loss of RAGE expression, likely indicative of a decline in AT1 cells, with a corresponding increase in PanCK+ cells, representing cancerous cells (**Figure 3d**). The PanCK+SFTPC+ AT2 cells, which are believed to be the cells of origin for LUAD, remained relatively stable, displaying only a slight decrease throughout progression.

The lack of a substantial increase in tumor cell numbers during later stages prompted us to revisit our multiplexed images to investigate what might be happening with the tumor cells, seeking to resolve this unexpected observation.

Through examining both the tumor and stromal microenvironments, we found compelling evidence supporting EMT to varying degrees. We observed epithelial cells beginning to express basal vimentin, a marker of EMT (**Figure 3g**, first row). Additionally, some epithelial cells appeared rounder, likely due to the loss of polarity, and were often detached from its original epithelium, suggesting they had gained mobility (**Figure 3g**, second row). In other cases, we observed groups of epithelial cells detaching collectively (**Figure 3g**, third row), with the outer layer of cells expressing vimentin as they floated in the luminal space. We confirmed these qualitative observations quantitatively by noting shifts in the proportions of epithelial and mesenchymal cells (**Figure 3a,d**) and the reciprocal expression of PanCK and Vim across different stages of disease progression (**Figure 3h**).

### Microenvironmental adaptations reflect the hallmark dynamics of LUAD progression

Following our independent analysis of the progression dynamics of tumor, stromal, and immune cells in LUAD (lung adenocarcinoma), we next sought to investigate the interactions among these cell types within the tumor microenvironment. To achieve this, we employed the UTAG^28^ algorithm to identify and characterize distinct microenvironments present in the dataset (**Figure 4a-b**).

Our analysis of the spatial distribution of tumor, stromal, and immune microenvironments revealed trends consistent with those observed at the cellular level. Specifically, the area occupied by epithelial or tumor microenvironments exhibited a significant increase from adjacent-normal tissues to tumor, with no substantial changes observed between different tumor subtypes (**Figure 4c**). In contrast, a significant expansion in the tumor-stroma interface area was noted, indicating pronounced stromal proliferation during tumor progression.

Further investigation into the spatial localization of immune cells revealed that a considerable proportion of lymphocytes originated from tertiary lymphoid structures (TLS) (**Figure 4c**)^29–31^. The area occupied by TLS was found to be larger in invasive adenocarcinomas, and this increase was associated with a progressive suppression of immune activity (**Figure 4d**). We observed a reduction in immune activation markers such as ICOS and GZMB. Concomitantly, an increase in the expression of immunosuppressive markers, including TIM3, was observed within macrophage networks, further contributing to immune suppression (**Figure 4e**).

To assess whether specific spatial rearrangements occurred during tumor progression, we compared the interactions between adjacent normal tissue and tumor samples. Notably, epithelial-like cells exhibited enhanced interactions both within epithelial-like cells and between epithelial-like cells and fibroblasts, compared to normal tissue (**Figure 4f**, black and blue dotted lines). In contrast, interactions between epithelial-like cells and immune cells were consistently reduced across all tumor subgroups (**Figure 4f**, blue dotted line), relative to normal tissues. These observations highlight three key spatial patterns: (1) the aggregation of tumor cells, (2) the encapsulation of tumor cells by fibroblasts, and (3) the formation of a fibroblast barrier surrounding tumor cells, which likely serves to impede immune cell infiltration (**Figure 4f**).

### LUAD progresses either through immunomodulation or fibrosis in patients

Lastly, we sought to explore whether the cellular patterns could be stratified at the patient level. In order to achieve this goal, we performed hierarchical clustering based on cellular densities derived from both imaging panels (**Figure 5a**). In total, we identified four distinct lesion groups: G1. In-situ lesion (32.8%, 40/122), G2. Macrophage-enriched population (5.7%, 7/122), G3. Inflammatory lesion (40.8%, 50/122), and G4. Fibrotic lesion (20.4%, 25/122).

**Figure 5:**
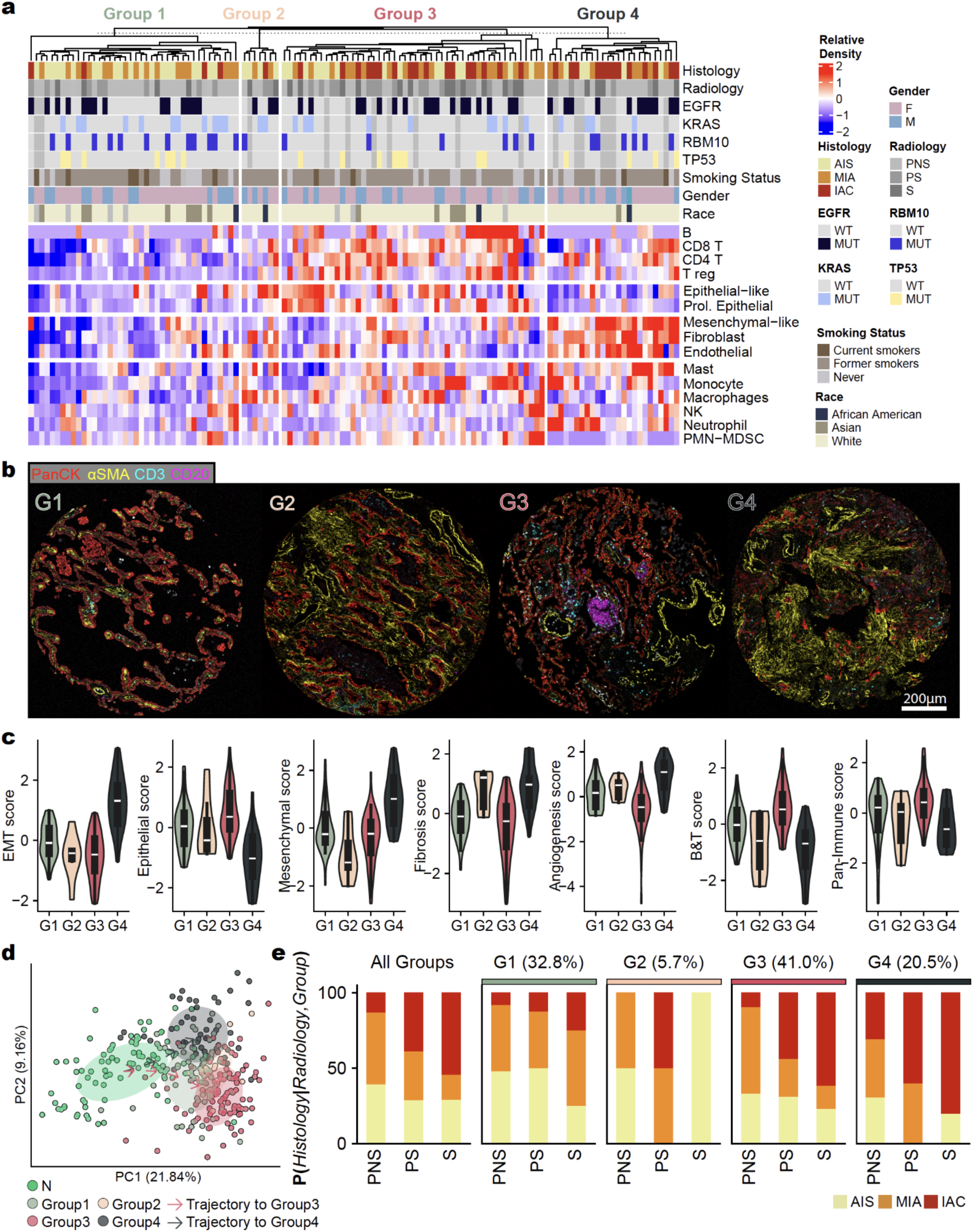
Patient stratification based on the cellular architecture of LUAD lesions. **a**) Heatmap visualizing cell type densityl for 122 patients. Hierarchical clustering was applied on B and T cells to categorize patients into 4 groups based on immune distribution. **b**) Representative PNS AIS images for each patient subgroup to demonstrate their immune hotness and fibrosis. **c**) Violin plot showing the EMT, fibrosis, angiogenesis, B cell and T cell infiltration, and general immune infiltration score. **d**) PCA plot from Figure 1e overlaid with IMC based patient groups. Ellipses in the figure represent the mean and the covariance of each subgroup in PCA space. Arrows depict trajectory from normal to Group 3 and Group 4. **e**) Conditional probability of developing an invasive adenocarcinoma given radiology and patient subgroup. Group 4 patients have 24.95% more chance of developing an invasive adenocarcinoma compared to all patients combined.

Group 1, the in-situ lesions, displayed relatively low density of most cell types, including epithelial and immune cells, indicating the mildest form of LUAD. The majority of these patients were classified as AIS (50.0%) or PNS (62.5%), with their histological images showing minimal signs of disease progression (**Figure 5b**, G1). Group 2, a minor population, comprised seven patients showing no advanced signs of disease progression. Group 3 was characterized by a high density of epithelial cells and infiltrating lymphocytes (**Figure 5b**, G3) with a more advanced disease state as indicated by histological (15 AIS, 18 MIA, 17 IAC, 42.5% IAC) and radiological (21 PNS, 16 PS, 13 S, 32.5% S) assessments. Group 4 exhibited significant fibrosis with comparatively lower immune infiltration (**Figure 5b**, G4).

Upon further examination of biological processes underlying Group 4, this lesions were exposed to most EMT, fibrosis, and angiogenesis (**Figure 5c**) relative to other groups. At the same time, this group also had the lowest number of immune cells (**Figure 5c**), likely exposing the patients to nutrition-rich invasive tumors that grow uninterruptedly. Histologically, these patients were predominantly diagnosed as aggressive and invasive (5 AIS, 7 MIA, 13 IAC), while radiologically, most were classified as mild (13 PNS, 5 PS, 5 S).

Considering the divergent nature of biological processes in Group 3 and Group 4 lesions (**Figure 5c**), we conjectured that LUAD progression trends would follow a non-linear binary trajectory. In order to test this idea, we went back to the PCA analysis from Figure 1e and overlaid the PCA plot with the identified groupings (**Figure 5d**). To identify the central tendency and dispersion of each group with respect to others, we added ellipses depicting the mean and covariance in the PCA plot. Based on the localization of the ellipses, we were able to separate normal, Group 3, and Group 4 ROIs as extremes and Group 1 and 2 as intermediates leading to either extreme.

Lastly, we aimed to quantify the risk associated with patients at the extremities. To achieve this goal, we calculated the conditional probability to evaluate the likelihood of disease invasiveness based on radiological assessment (**Figure 5e**). This analysis indicated that Group 4 patients with fibrotic lesions were up to 24.95% more likely to be diagnosed with IAC than the average patient, highlighting the necessity for heightened scrutiny during GGO-based diagnosis. Altogether, we report that fibrotic tumors, which are stromally enriched, were identified in 20.4% of all LUAD cases. These patients were frequently diagnosed as PNS through radiology but were identified as IAC through histology. This finding suggests that patients with fibrotic tumors may face an increased risk under current diagnostic procedures, which primarily determine whether a patient should undergo biopsies based on radiological assessments.

## Discussion

In this study, we presented a comprehensive spatiomolecular atlas of early-stage LUAD progression at single-cell resolution. Because early-stage LUAD patients are rare, representing only 25% LUAD patients^32,33^, this collection of 122 patients with paired adjacent-normal samples enables characterization of LUAD progression at unprecedented scale and resolution. Through IMC, we showcase an overview of LUAD progression that is exceptionally valuable due to its rarity, specificity, and relevancy.

Our analysis delineated the shifts in molecular landscape of LUAD lesions by histological and radiological subtypes, enabling the separation between cellular changes exclusive to tumor initiation (**Figure 2-3**) with those that intensify as the disease progresses. This temporal differentiation not only provides clear data-driven evidence, but also aligns with existing knowledge of cellular roles: innate immune cells offer immediate defense with limited long-term impact, while adaptive immune cells are equipped with mechanisms that evolve to counteract cancerous mutations effectively.

We revealed nuanced modulation of immune functions along the tumor progression axis, marked by the inhibition of immune activation molecules and the concurrent activation of suppressive functions, irrespective of their initial abundance. Adaptive immune cells were often found in the form of TLS, enhancing T cell recruitment and infiltration. We also shared evidence supporting the simultaneous cytotoxicity and regulation, likely giving rise to the prevalent inflammation^34–36^ within the tumor immune microenvironment.

Given the substantial presence of T cells in the TME, patients with predominantly inflammatory lesions could benefit from targeted immunotherapies. Specifically, the use of anti-CTLA-4 immunotherapy^37–41^, possibly in conjunction with anti-ICOS therapies^42–44^, could reactivate the anti-tumor immunity mediated by TLS, offering a promising strategy to enhance treatment efficacy.

Our findings also underscored the role of fibrosis in the development of LUAD^45^. The presence of fibroblast has been increasingly recognized as a marker of tumor aggressiveness^46^, influencing tumor behavior through mechanisms such as the activation of growth factors, inflammatory immunomodulation, induction of hypoxia, and exertion of oxidative stress. We observed that fibroblasts form an insulating layer around the basal membrane of tumor cells, effectively shielding them from direct immune contact (**Figure 4f**). At the patient level, this phenomenon correlated with an overall reduction in immune activity (**Figure 5c**, G3), differentiating the evolutionary pattern of fibrotic LUADs with inflammatory LUADs.

Given the critical role of fibroblasts in promoting tumor progression and resistance to treatment, targeting pro-fibrotic signals from tumors presents a promising therapeutic avenue. Several emerging therapies aim to modulate fibroblast activity or reduce their presence within tumors. For instance, drugs that inhibit the signaling pathways activated by fibroblasts, such as TGF-β inhibitors, have shown potential in reducing fibrosis and enhancing the efficacy of existing cancer treatments. Additionally, therapies that directly target fibroblast-to-cancer cell signaling could disrupt the protective environment fibroblasts create around tumor cells, thereby enhancing immune system access and activity within the TME. Clinical trials exploring agents that target fibroblast activation proteins, which are highly expressed in cancer-associated fibroblasts, are also underway. These therapies not only aim to diminish the fibrotic barrier but also seek to prevent the immunosuppressive effects mediated by fibroblasts, offering a dual approach to combating tumor growth and invasiveness. As research progresses, the integration of fibroblast-targeting strategies into standard cancer treatment regimens could significantly improve outcomes for patients with fibrotic LUAD.

Clinically, our findings suggest a need to reevaluate how traditional clinical assessments using these modalities are understood^46–50^. Our study reveals that while current radiological techniques are capable of accurately identifying the majority of the patients (**Figure 5a**, Group 1 and Group 4), accounting for 73.8% of our cohort, they fail to specifically assess the risk for 20.5% of patients. Given that radiological assessments are typically the first line of diagnostic inquiry prior to more invasive procedures, there is a critical need to enhance the detection and management of such fibrotic tumors at a non-invasive level. These findings underscore the importance of heightened vigilance and specialized care in the evaluation of such cases to better inform treatment strategies and improve patient outcomes.

Nevertheless, there were several noteworthy limitations to the study design. The first limitation of the study was that the assessment of risk, commonly achieved through survival rate, was challenging. Tracking the clinical outcomes for most patients was impractical as more than 80% of early-stage localized LUAD patients are considered disease-free post-surgery. To overcome this issue, we took the invasiveness criteria in histopathological assessment as an anchor point to determine the risk of each group identified by IMC molecular subtype (**Figure 5e**).

The 1mm^2^ restricted field of view from IMC constrained the capability to capture the entirety of GGO nodules, which typically range from 1-2 cm in diameter. Another limitation was the lack of longitudinal disease progression samples due to the patient-driven nature of the study. To a certain extent, the large cohort size mitigates concerns about sampling coverage and the lack of longitudinal samples. In total, this study examined 2.24 million cells across 701 ROIs spanning a tissue area of 742.17 mm² from 122 patients. We believe that the extensive cohort size compensates for the concerns towards the lack of sample coverage.

In conclusion, this study presented the first deep spatiomolecular profiling of early-stage LUAD, utilizing single-cell resolution IMC to explore the complex interplay of cellular phenotypes within the TME. Our comprehensive analysis not only underscores the molecular heterogeneity prevalent in early-stage LUAD but also highlights critical transitions in cellular behavior from tumor initiation to progression. The findings emphasize the limitations of current diagnostic modalities, advocating for an integrated approach that combines traditional imaging and histopathology with advanced molecular profiling techniques to enhance diagnostic precision and therapeutic targeting.

The data derived from this study lay a groundwork for redefining patient stratification and treatment planning, incorporating detailed molecular insights that could lead to more personalized therapy approaches. This could particularly impact the management of patients with fibrotic tumors, where targeted therapies might significantly improve outcomes. As the field moves forward, the integration of such detailed molecular data with clinical practice promises to refine our understanding of LUAD, ultimately leading to better patient outcomes and a deeper understanding of the disease’s underlying biology.

## Methods

### Imaging Mass Cytometry TMA staining

A total of 20 unstained TMA slides were shipped to SironaDx for IMC staining. Two different panels with a total of 72 antibodies were stained for this study. Most of the antibodies used in both the panels were custom conjugated using the MaxPar X8 multimetal labeling kit (Standard BioTools) according to the manufacturer’s protocol^51^. All the antibodies were tested on relevant control tissues such as tonsil, lymph node, Invasive squamous cell carcinoma and DLS lung tissues with TLS and immune cell infiltrates. The staining quality and specificity were verified by in-house pathologists at SironaDx. All the TMA slides were stained following the standard protocol prescribed by Standard BioTools. Briefly, freshly cut 5μm TMA slides were pre-warmed and baked at 650C for 2 hours. The slides were deparaffinized by dipping in CitriSolv solution 2X for 15 mins each (RT) followed by rehydration in series of 100%, 100%, 80% and 75% ethanol, 5 mins each at RT. After thorough wash in MilliQ water, the slides were dipped in basic AR solution at 960C for 30 mins. The slides were then cooled to room temperature and washed with TBS and MilliQ water. The slides were blocked in ThermoFischer Scientific SuperBlock solution (Cat No: 37515) for an hour while the antibody cocktail was prepped for staining. The slides were incubated with the antibody cocktail overnight in 40C and washed with 0.2% Triton X-100 in PBS (2X) followed by 2X TBS wash. Cell nuclei were stained with Intercalator-Iridium diluted in PBS for 30 mins at RT. Slides were briefly washed with MilliQ purified water and air dried prior to ablation.

### Preprocessing IMC data

We used the IMC package (https://github.com/ElementoLab/imc, version 0.1.4) to preprocess the raw IMC data into a combined AnnData object that contains the location and expression profiles of acquired cells. With original MCD files acquired through Hyperion machine, ROIs were extracted as stacks of images in tiff file format along with associated metadata including channel and epitope information. To identify cellular and non-cellular region from the images, we used a pretrained ilastik^52^ model to predict nuclear, cytoplasm, and background probability. The probability mask was subsequently segmented using DeepCell^53^ to capture and identify cellular and nuclear borders.

To quantify cellular expressions, we used the cell masks to aggregate the mean intensity of pixels within a cell for each antibody channel through scikit-image. We combined the per cells expression vector from all cells across all images into a single matrix through scanpy^54^ in anndata^55^ format to process the data comprehensively and consistently. We then performed log transformation, Z-score normalization with truncated at positive and negative 3 standard deviations, followed by harmony^56^ (version 0.3.0) data integration to phase out sample-specific biases.

### Cell type identification

To systematically and quantitatively identify the cell phenotypes captured in IMC with all markers combined across all images, we performed PCA, neighborhood calculation using 15 nearest neighbors, and Leiden clustering^57^ at resolution 0.5 to the normalized and batch corrected cell expression. These clusters were broadly annotated into epithelial, stromal, and immune cell groups based on their expression. We repeated the clustering procedure for each of these broad cell types for a more granular subgroup of cells.

### Identification and comparison of TME

To capture adjacent normal structures and TME, we applied UTAG^28^ across the full dataset with a *min_dist* of 20 micrometers and used K-means clustering to create 20 clusters for niche identification. Functional analysis across GGO conditions were performed using scanpy rank_genes_groups function, where each condition was statistically compared using t-test with all other conditions combined. All resulting comparisons were Benjamini-Hochberg corrected. Resulting difference difference in marker enrichment was visualized as dotplots (**Figure 4d-e**).

### Statistical comparison of cellular and microenvironment composition

To quantify the difference in cellular densities across GGOs, each ROI was aggregated by the number of cells normalized by area. We performed a pairwise two-sided Mann-Whitney U test between all conditions using pingouin^58^ (version 0.3.12). These p-value significances were Bonferroni-Hochberg corrected to adjust for the generated multiple hypothesis (**Figure 2a-c, f-g, j-k, 3a-b, 4c, Extended Figure 4**). The cellular composition of identified microenvironments explain the physiological role of each microenvironment (**Extended Figure 5**)

### Principal component analysis on IMC panels

To look at general trends as well as tumor heterogeneity of our samples, we integrated our dataset as a single numerical matrix where each data point was the cellular density of an ROI. For each ROIs, the two matrices representing each panel were concatenated column-wise. We standardized the impact of each density by z-score to capture a more diverse cell type portfolio. We performed PCA on the scaled data and plotted the first two principal components overlayed with the histopathology grading on **Figure 1e** using seaborn scatterplot. Individual images with RGB colors were generated with the MCD viewer software from Standard Biotools.

### Quantification of cellular interactions

To quantify cellular interactions for each GGO condition without cellular density biases, we calculate interaction frequencies normalized by expected interaction for a given cellular densities. The observed interaction frequency was calculated by counting the pairwise interaction between cell types in a spatial network where cells were defined as connected if within 100 µm radius. The expected interaction frequency was calculated by randomly shuffling cell identities 200 iterations. We add one to both the observed interaction frequency and expected frequency to avoid division by zero. Log-fold change for cell type interaction was then computed by taking differences between two log-transformed values. Since each of these log-fold changes in interactions were calculated on a per-ROI level, we aggregated these for each GGO condition by taking the mean log-fold change for each condition. Differences in cellular interaction across conditions were measured by subtraction of the means.

### Hierarchical clustering of patients

To stratify patients based on high-information content IMC cellular characteristics, we combined the cell densities captured from both the tumor and immune panels at the patient level. First, we aggregated the total number of cells per cell phenotype and the total area covered by all tumor ROIs. Then, for each panel, we recalculated the overall density of cells for each cell type. To integrate the two panels, we concatenated the unique cell phenotypes and calculated the mean density for overlapping cell types. Finally, we performed hierarchical clustering through the hclust function provided by the ComplexHeatmap package (v 2.18.0) to robustly generate four patient subgroups (**Figure 3a**).

### Assessment of risk for patient subgroups

We performed scoring and conditional risk assessment to elucidate the meaning and consequences each patient subgroup was subject to (**Figure 5c,e**). In order to score the biological processes each patient was exposed to, we used the scores_genes function from scanpy tools. For the assessment of patient risk, we calculated the observed frequency of histological subtypes for each radiology and patient subgroups. The subsequent conditional probability was calculated by normalizing the observed frequency for radiology and patient subgroups jointly.

## Data availability

The raw and processed dataset is publicly available at the following URLs:

- Raw data: https://zenodo.org/records/14822106
- Processed spatial data: https://zenodo.org/records/14827075

## Code availability

Source code is publicly available at the following URL: https://github.com/ElementoLab/ggo-imc

## Acknowledgements

The project has been funded by Johnson and Johnson. O.E. is supported by NIH grants UL1TR002384, R01CA194547, and Leukemia and Lymphoma Society SCOR 7012-16, SCOR 7021-20 and SCOR 180078-02 grants. This work was also supported in part by NIH grant UH3CA244697, and funds from The Neuberger Berman Foundation Lung Cancer Research Center. A.F.R. is supported by Angelini Ventures S.p.A. Rome, Italy.

## Author contributions

J.K., H.R., B.B, L.Y., A.S., S.B., E.H., V.M., T.M., A.F.R., and O.E. planned the study; A.S. coordinated the clinical sample acquisition and biobanking. H.R. stained the IMC TMA. J.K. performed computational analysis. T.M., A.F.R., O.E. supervised the research. J.K. and K.F. wrote the manuscript. B.B., L.Y, T.M., A.F.R., O.E. edited and provided feedback to the manuscript.

## Extended data figures

**Extended Figure 1:**
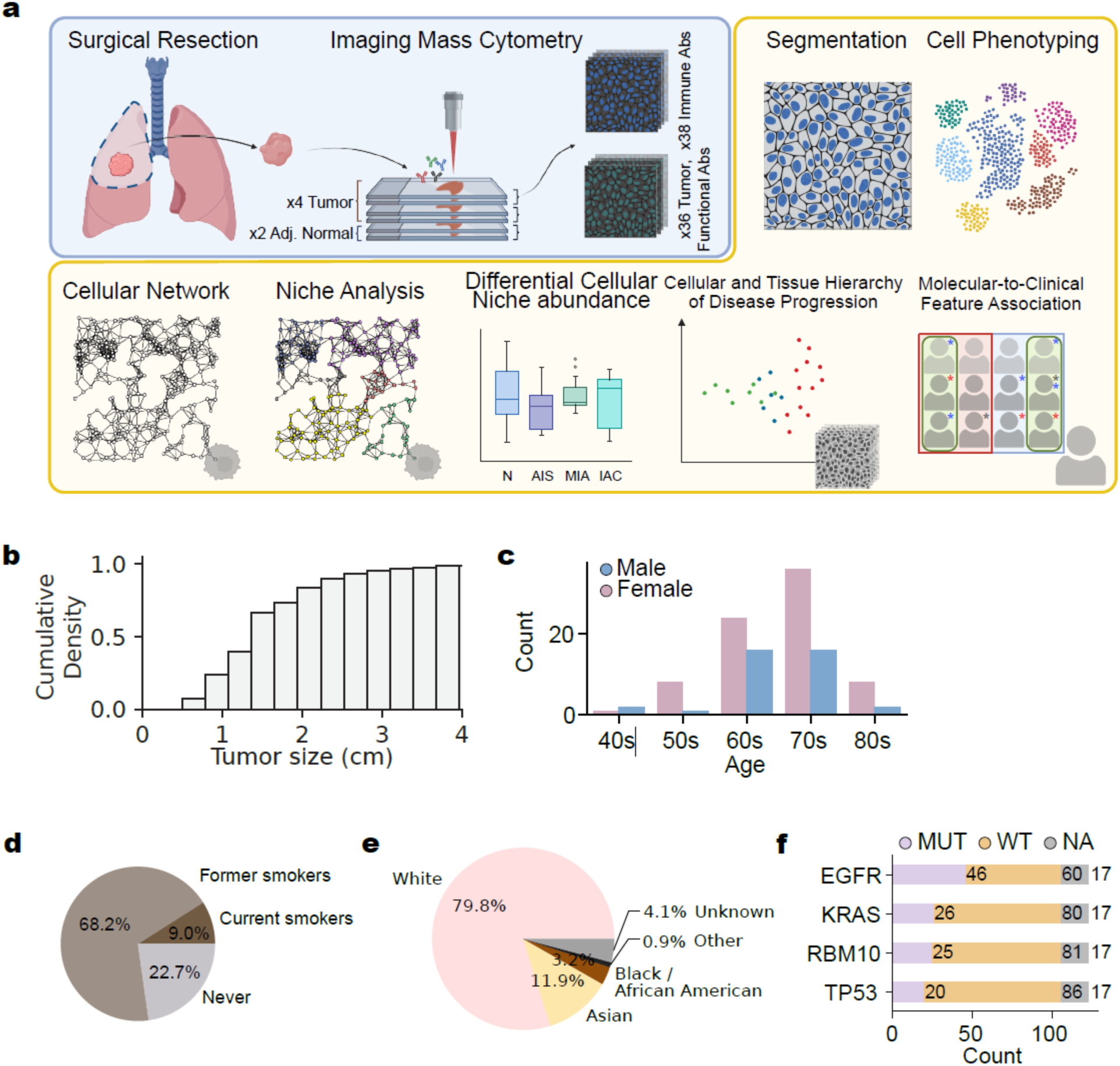
Study design and patient demographics. **a**) Schematic of the study design broken down to blue box representing sample collection and orange box depicting computational analysis. Every patient in the study had their lung surgically resected. We scanned two adjacent stack semi-replicates tissue slides for two sets of representative tumor regions and one adjacent normal, collecting in total up to 6 tissue slides per patients. On the computational side, cells were segmented, phenotyped, and analyzed. **b**) Distribution of tumor size. The cohort came from early-stage LUAD patients, and thus were mostly less than 2cm in size. **c**) Patient gender and age distribution. 66% of patients were female. 90% of patients were of age greater than 59. **d**) Patient smoking history visualized through a pie chart. 77.3% of patients had a history of smoking. **e**) Patient race distribution. While majority of patients were white, the cohort also captures a considerable amount of Asian and African American population. **f**) Visualization of four most common mutation detected through WES.

**Extended Figure 2:**
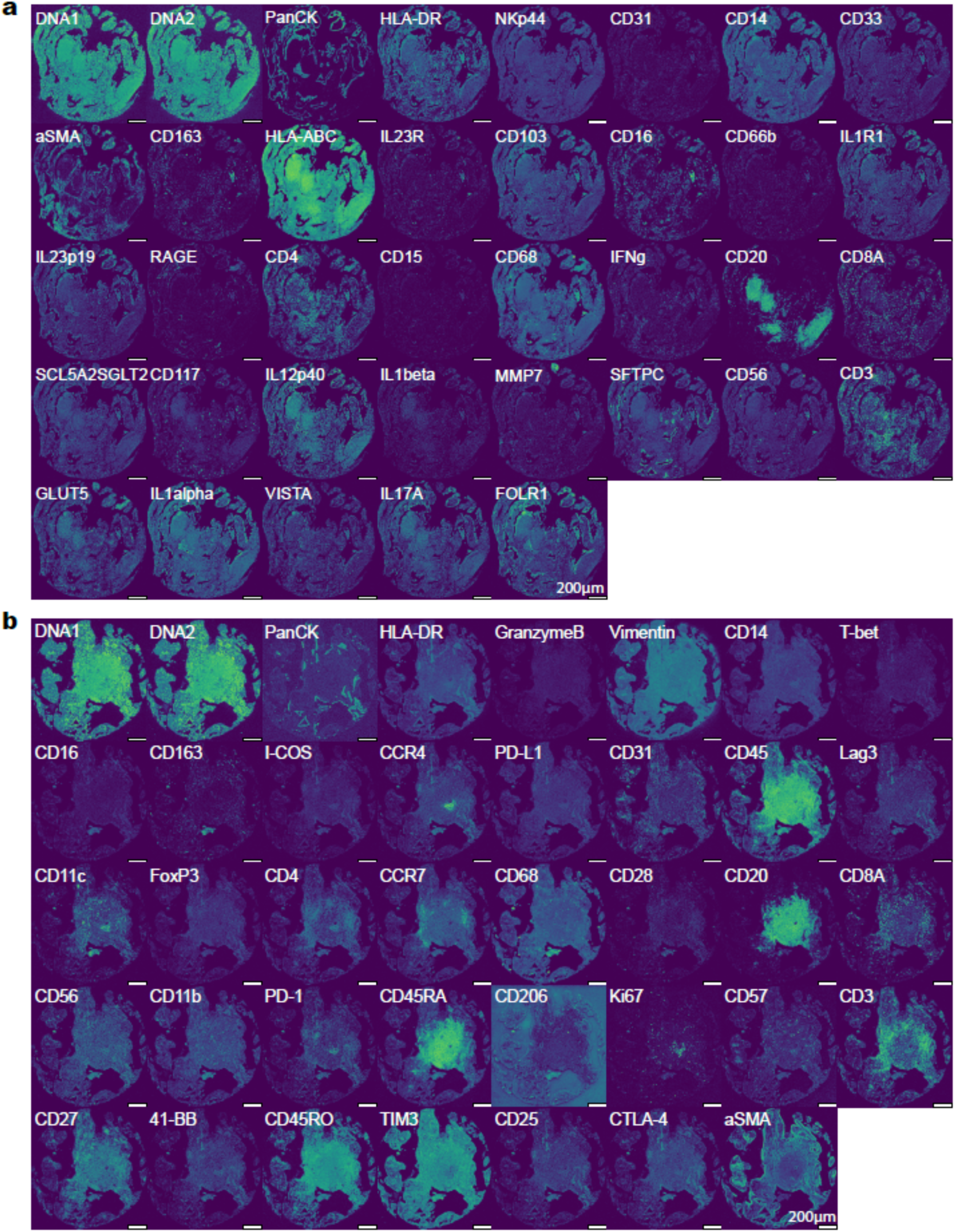
Illustration of collected sample IMC data. Illustration visualizing the two panels of adjacent z-stack ROI. **a**) Visualization of the tumor panel. **b**) Visualization immune panel.

**Extended Figure 3:**
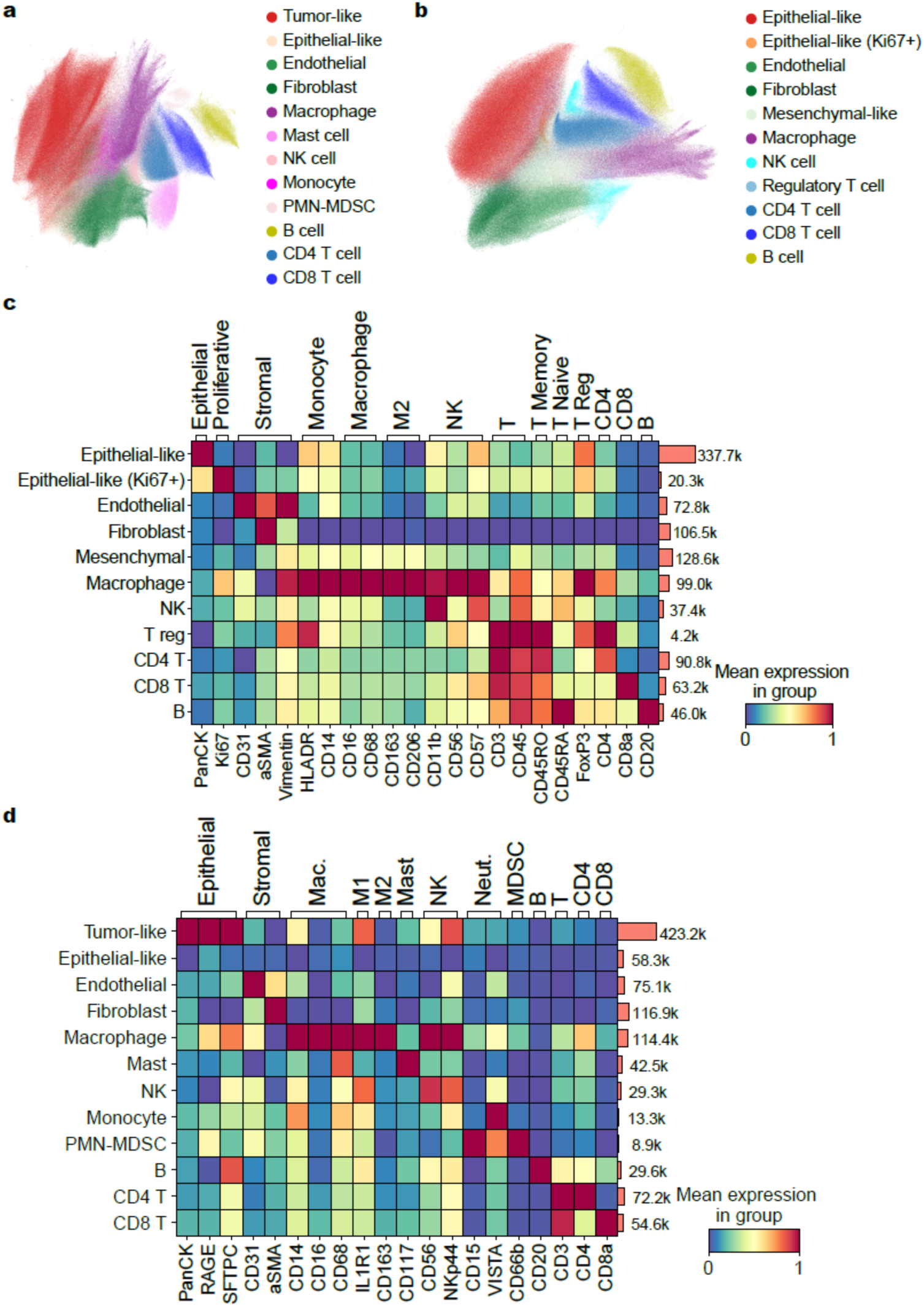
UMAP and expression profile of captured cells. **a**) Violin plot showing the distribution of tumor sizes stratified by GGO solidity. **b**) Clinical tumor size across radiological subtypes of LUAD. * p<0.05, ** p<0.01, **** p<0.0001, two-sided Mann-Whitney U-test with BH multiple hypothesis correction. **a**) UMAP of cell types from tumor tailored panel. **b**) UMAP of cell types from immune tailored panel. **c**) Heatmap of mean expression profiles for cell types in immune tailored panel. Min-max scaled on a per column basis. Bar plot with numbers on the right indicates the number of cells within each group. **d**) Heatmap of mean expression profiles for cell types in tumor tailored panel. Min-max scaled on a per column basis. Bar plot with numbers on the right indicates the number of cells within each group.

**Extended Figure 4:**
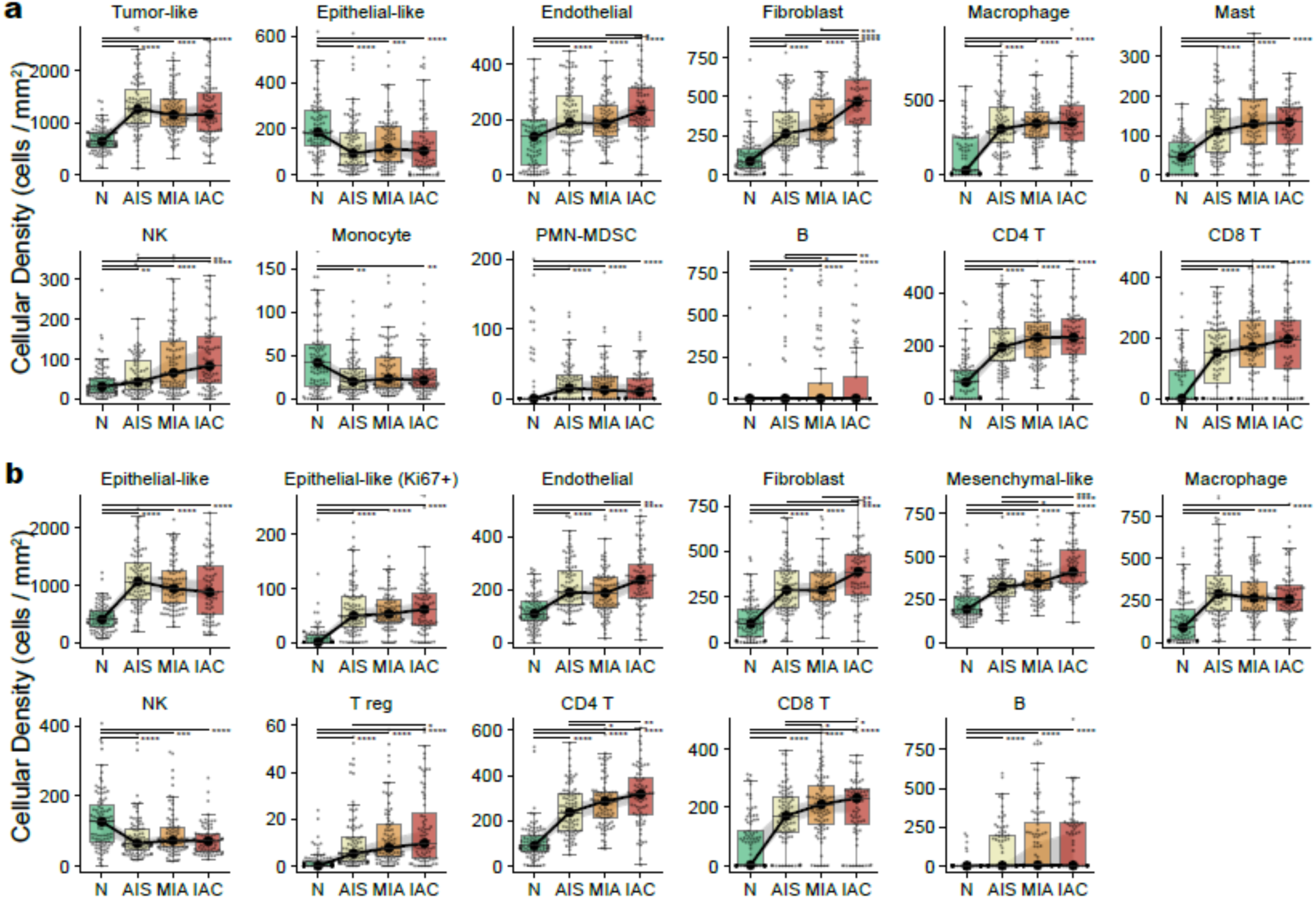
Cellular abundance characterized across patients. **a-b**) Statistical comparison of cellular densities across cell types. Extent of disease onset and progression for each sample was identified for histopathology. **a**) visualize cell type density for the immune-tailored panel and **b**) illustrate cell type density for the tumor-tailored panel. Bars and stars on top of box plots represent statistical significance between pairwise densities across conditions. * p<0.05, ** p<0.01, *** p<0.001, **** p<0.0001, two-sided Mann-Whitney U-test with BH multiple hypothesis correction.

**Extended Figure 5:**
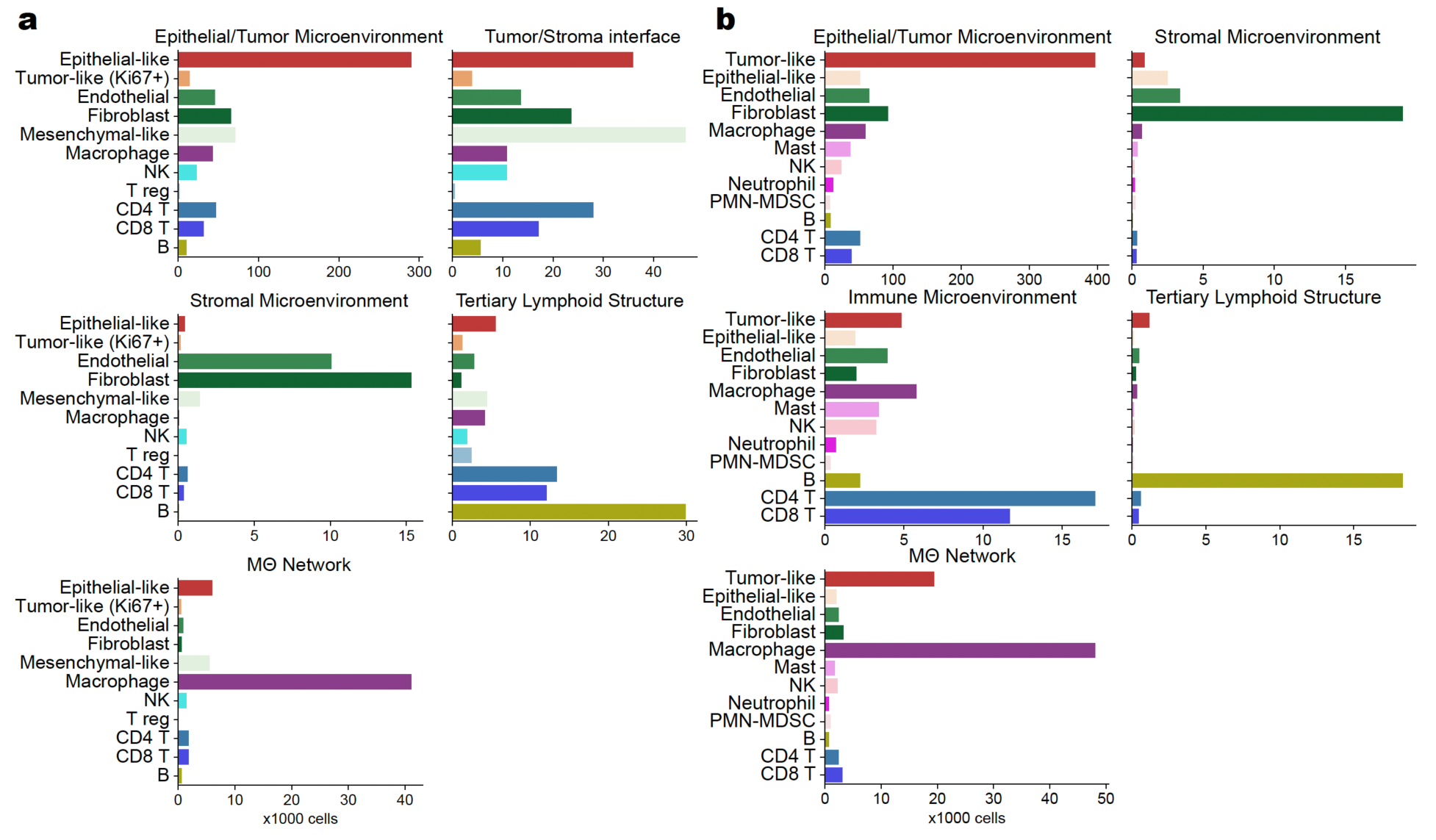
Cellular characterization of niches and their functions. **a**) Cellular composition of detected niches for the immune tailored panel. **b**) Cellular composition of detected niches for the tumor tailored panel. **c**) Hallmark pathway analysis on epithelial cells from external single cell RNA sequencing data.

